# PyVar: An Extensible Framework for Variant Annotator Comparison

**DOI:** 10.1101/078386

**Authors:** Julie Wertz, Qianli Liao, Thomas B Bair, Michael S Chimenti

## Abstract

Modern genomics projects are generating millions of variant calls that must be annotated for predicted functional consequences at the level of gene expression and protein function. Many of these variants are of interest owing to their potential clinical significance. Unfortunately, state-of-the-art methods do not always agree on downstream effects for any given variant. Here we present a readily extensible python framework (PyVar) for comparing the output of variant annotator methods in order to aid the research community in quickly assessing differences between methods and benchmarking new methods as they are developed. We also apply our framework to assess the annotation performance of ANNOVAR, VEP, and SnpEff when annotating 81 million variants from the ‘1000 Genomes Project’ against both RefSeq and Ensembl human transcript sets.

## Introduction

High-throughput sequencing pipelines for genomic research and clinical use generally incorporate a downstream step to integrate predictions about the functional consequences of variation against a reference sequence at coding and non-coding sites in an individual’s genome. This procedure relies first on accurate calling of the single-nucleotide polymorphisms (SNPs) by specialized methods [1] and second, on downstream assignment of consequences to those SNPs, so-called ‘variant annotation.’

A major challenge in accurate variant annotation is the apparent lack of concordance when annotating against different transcript sets or between different annotation algorithms. Others have shown that the use of different transcript sets as a basis for annotation can dramatically affect the outcome of the annotation calls [2]. Additionally, even if the same transcript set is used between methods, the method must select one or several of the potentially many transcripts to annotate against. This leads to differing outcomes in annotations which may arise from different logic structures in the algorithms or different user criteria for annotation. Unfortunately, incorrect annotations or disagreement in annotation outcomes can lead investigators to waste resources tracking down variants of little interest or to miss severe variants of potential clinical significance.

There is a need for comparison of the many competing methods for variant annotation and for easy, automated benchmarking of constantly changing existing methods as well as newly developed methods. Here, we present an easily extensible python framework, PyVar, for automated analysis of variant annotator methods on a common dataset. PyVar allows users to quickly integrate the output of new or updated variant annotation methods into its benchmarking workflow by providing simple class constructors to standardize the output of different methods. The framework automatically produces rich HTML graphics for exploratory analysis. A unique feature of PyVar is that we have attempted to standardize the annotation consequences from each annotator into a common ontology, simplifying downstream analysis. Heatmaps of lognormalized, most severe standardized and non-standardized consequences are created automatically. We used our PyVar framework to analyze 81 million SNPs from a publically available repository at the 1000 Genomes Project [3] with three popular variant annotator methods: ANNOVAR [4], SnpEff [5], and VEP [6]. We will discuss a comparison of the results for these three methods and novel insights into the cause of discrepancies between methods.

## Results

### The PyVar framework

PyVar uses a set of custom python classes to translate from the disparate output formats and ontologies of the various annotator methods into a common format and ontology. Included in the code are classes for analysis of various ANNOVAR, VEP, and SnpEff output formats. Each class inherits from a general ‘AnnotationFile’ superclass, making the code easily extensible to other formats and annotators for quick validation and testing of new methods.

The translation tables to convert from the output ontologies of ANNOVAR, SnpEff, and VEP can be found in Table 1. Some compromises were necessary to reclassify each type of consequence under a common ontology in order to simplify the comparisons. For example, VEP’s classifications of ‘splice_donor_variant’, ‘splice_acceptor_variant’, and ‘splice_region_variant’ were all reclassified to ‘splicing’ because ANNOVAR doesn’t have an equivalent consequence assignment. Similarly, for ANNOVAR, the ‘stoploss’ and ‘stoploss SNV’ terms were simplified to ‘stoploss.’ In a few cases, the reclassification was arbitrary (e.g., ANNOVAR’s ‘upstream;downstream’ classification was reclassified to ‘upstream’).

**Table 1.** Standardized consequences translation table.

| PyVar | ANNOVAR | VEP/SnpEff |
| --- | --- | --- |
| **frameshift** | frameshift deletion | frameshift_variant |
| **frameshift** | frameshift insertion | — |
| **frameshift** | frameshift_block_substitution | — |
| **stopgain** | stopgain | stop_gained |
| **stopgain** | stopgain SNV | — |
| **stoploss** | stoploss | stop_lost |
| **stoploss** | stoploss SNV | — |
| **splicing** | splicing | splice_donor_variant |
| **splicing** | exonic;splicing | splice_acceptor_variant |
| **splicing** | — | splice_region_variant |
| **inframe_deletion** | nonframeshift deletion | inframe_deletion |
| **inframe_insertion** | nonframeshift insertion | inframe_insertion |
| **inframe_insertion** | — | disruptive_inframe_insertion |
| **inframe_deletion** | — | disruptive_inframe_deletion |
| **synonymous** | synonymous SNV | synonymous_variant |
| **synonymous** | — | stop_retained_variant |
| **nonsynonymous** | nonsynonymous SNV | start_lost |
| **nonsynonymous** | nonframeshift_block_substitution | initiator_codon_variant |
| **nonsynonymous** | — | missense_variant |
| **nonsynonymous** | — | incomplete_terminal_codon_variant |
| **UTR3** | UTR3 | 3_prime_UTR_variant |
| **UTR5** | UTR5 | 5_prime_UTR_variant |
| **UTR5** | UTR5;UTR3 | 5_prime_UTR_premature_start_codon_gain_variant |
| **upstream** | upstream | upstream_gene_variant |
| **upstream** | upstream;downstream | — |
| **downstream** | downstream | downstream_gene_variant |
| **intron** | intronic | intron_variant |
| **intergenic** | intergenic | intergenic_variant |
| **intergenic** | — | intergenic_region |
| **nc_exon** | ncRNA_exonic | non_coding_transcript_exon_variant |
| **nc_exon** | — | non_coding_exon_variant |
| **nc_intron** | ncRNA_intronic | non_coding_transcript_variant |
| **splicing** | ncRNA_splicing | — |
| **splicing** | ncRNA_exonic;splicing | — |

Because all three methods can report multiple consequences for each variant (n.b., ANNOVAR does not by default, you must ask with the ‘—separate’ flag), a ranking of consequence severity had to be established to order the consequences in the heatmap plots (below) and to put only the most severe consequence for each annotator in the plot data. For the purposes of this work, we took the Ensembl/VEP severity ranking scale (http://www.ensembl.org/info/genome/variation/predicted_data.html#consequences) as our basis for most to least severe consequence:

‘frameshift’ > ‘stopgain’ > ‘stoploss’ > ‘splicing’ > ‘inframe_insertion’ > ‘inframe_deletion’ > ‘nonsynonymous’ > ‘synonymous’ > ‘UTR5’ > ‘UTR3’ > ‘nc_exon’ > ‘nc_intron’ > ‘intron’ > ‘upstream’ > ‘downstream’ > ‘intergenic’ > ‘None’

Note that ‘None’ is assigned by PyVar in certain circumstances and is not assigned by default by the three annotator methods. It is impossible to objectively rank consequences in severity for all potential situations. For example, it is possible to imagine situations where a ‘stop-gain’ near the end of a sequence may be less deleterious than a nonsynonymous missense variant at an important enzymatic region or binding site.

### Comparison of variant annotation methods on the 1000 Genomes callset

We compared the annotation outcomes between ANNOVAR, VEP, and SnpEff on the Ensembl and RefSeq transcript sets using PyVar. Comparison statistics are summarized in Table 2.

**Table 2.** Comparison of variant consequences for ANNOVAR, VEP, & SnpEff on 1000 Genomes Variant Set

|  | Method 1: ANNOVAR;<br>Method 2: VEP | Method 1: ANNOVAR;<br>Method 2: SnpEff | Method 1: SnpEff; Method 2:<br>VEP |
| --- | --- | --- | --- |
| Positional Match % | 99.7 | 99.7 | 100 |
| Transcript Match % (Ensembl) | 61.5 | 61.5 | 99.9 |
| Transcript Match % (RefSeq) | 44.7 | 50.2 | 85.8 |
| Most abundant consequences (with >= 1 shared Ensembl transcript between methods) | 'intronic' (61.8%) 'non-coding intron' (18.9%) 'downstream' (8.9%) 'upstream' (6.5%) | 'intron' (76.7%) 'downstream' (10.3%) 'upstream' (7.7%) | 'intron' (76.6%) 'downstream' (13.9%) 'upstream' (6.4%) |
| Most abundant consequences (with >= 1 shared RefSeq transcript between methods) | 'intronic' (78.8%) 'downstream' (5.9%) 'non-coding intron' (5.5%) 'upstream' (5.0%) | 'intron' (82.6%) 'downstream' (6.6%) 'upstream' (5.6%) | 'intron' (84.0%) 'downstream' (6.3%) 'upstream' (5.1%) |
| Most abundant consequences method 1 (no shared Ensembl transcripts) | 'intergenic' (61.3%), 'non-coding exon' (32.3%) | 'non-coding intron' (100%) | 'intergenic' (52.9%) 'non-coding exon' (29.1%) 'downstream' (10.6%) |
| Most abundant consequences method 2 (no shared Ensembl transcripts) | 'non-coding intron' (25.8%) 'nonsynonymous' (22.2%) 'downstream' (18.9%) 'upstream' (17%) 'synonymous' (10.9%) | 'intron' (99.9%) | 'non-coding intron' (66.0%) 'UTR5' (21.4%) 'intron' (3.7%) 'downstream' (3.3%) |
| Most abundant consequences method 1 (no shared RefSeq transcripts) | 'UTR3' (39.6%) 'intron' (20.3%) 'intergenic' (10.1%) 'UTR5' (8.8%) | 'non-coding intron' (100%) | 'UTR3' (38.4%) 'intronic' (27.9%) 'intergenic' (8.1%) 'UTR5' (6.8%) |
| Most abundant consequences method 2 (no shared RefSeq transcripts) | 'downstream' (41.0%) 'upstream' (21.8%) 'nonsynonymous' (13.5%) 'non-coding intron' (7.6%) | 'intron' (99.7%) | 'downstream' (50.6%) 'upstream' (23.0%) 'non-coding intronic' (6.1%) 'nonsynonymous' (5.8%) |

#### ANNOVAR versus VEP

ANNOVAR and VEP agreed on variant position in 99.7% of cases (84,587,501 variants). In 0.3% of cases, the methods didn’t agree on genomic position owing to discrepancies in the way that indels are reported in the output format of either method. These positions were excluded from downstream analysis.

However, when running ANNOVAR and VEP with Ensembl transcripts, there were significant differences in the specific transcript that each method chose to annotate across all of the variants (Figure 1). These differences led to differences in the assessment of the consequence of the SNP even though there was agreement on the position. For example, 38.5% of variants were annotated with completely different transcripts, whereas 61.5% shared at least one transcript selected for annotation between methods. For the RefSeq transcript set, 44.7% of variants were annotated with a common transcript.

**Figure 1.**
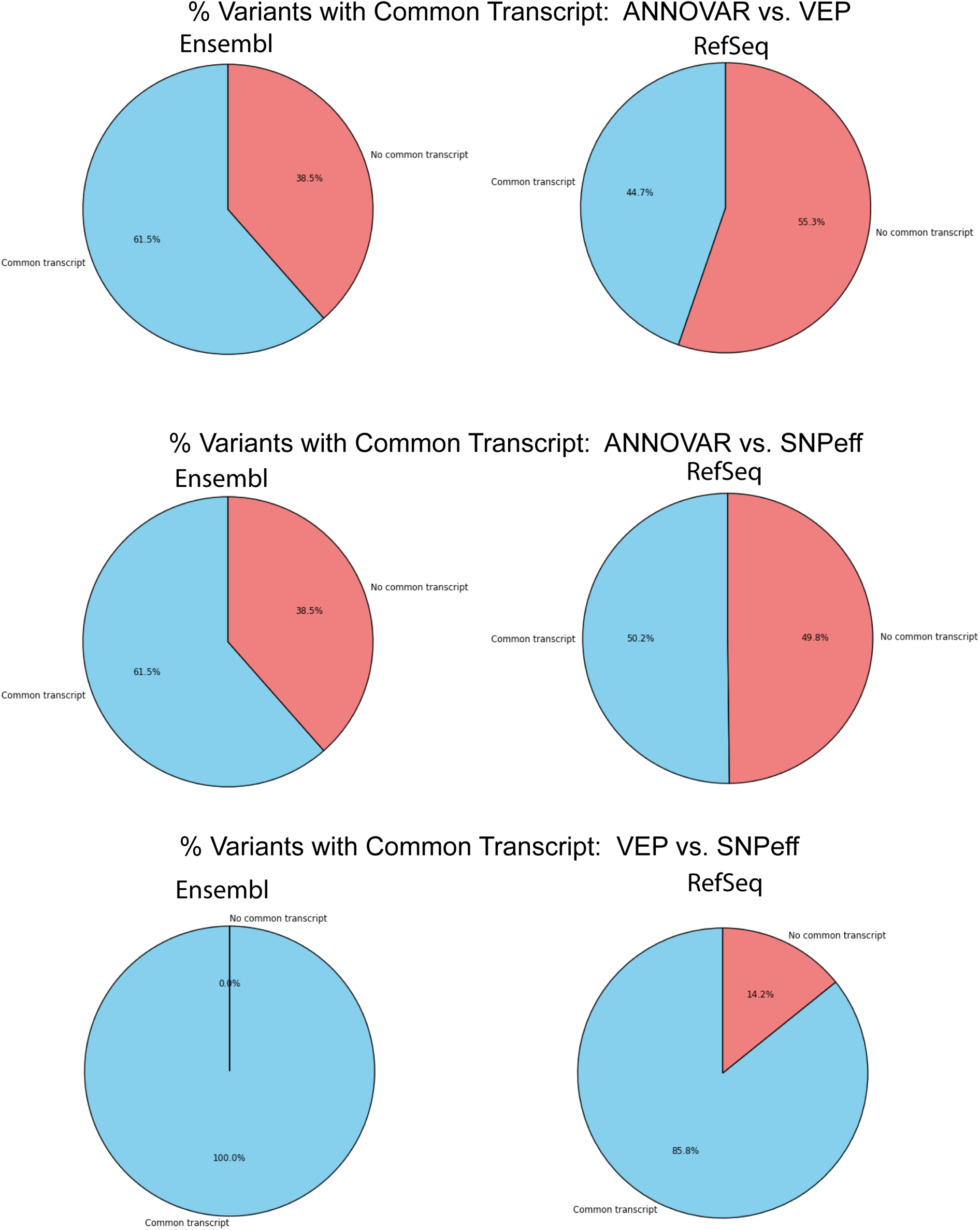
Percentage of variants with common transcript (a transcript that both annotators used for annotation). Variants with position mismatches are ignored.

Excluding the variants for which the methods selected non-overlapping transcripts to report, we looked at the overlap of normalized consequences among the remaining 61.5% (Ensembl) and 44.7% (RefSeq) of variants where a common transcript was reported. For both Ensembl and RefSeq, greater than 99% of variants had at least one consequence in common that was identified by both annotators. Of the variants sharing a common Ensembl transcript, ‘intronic’ (61.8%), ‘non-coding intron’ (18.9%), ‘downstream’ (8.9%), and ‘upstream’ (6.5%) were the most common annotations. Similarly, for RefSeq transcripts, ‘intronic’ (78.8%), ‘downstream’ (5.9%), ‘non-coding intron’ (5.5%), and ‘upstream’ (5.0%) were the top categories.

For the Ensembl transcript set, 1836 variants were annotated uniquely by ANNOVAR as ‘intergenic’ (61.3%) and ‘non-coding exon’ (32.3%). VEP annotations of the 3480 variants not shared with ANNOVAR included ‘non-coding intron’ (25.8%), ‘nonsynonymous’ (22.2%), ‘downstream’ (18.9%), ‘upstream’ (17%), and ‘synonymous’ (10.9%). With the RefSeq transcripts, a total of 27,306 variants were given discordant annotations by ANNOVAR, with ‘UTR3’ (39.6%), ‘intron’ (20.3%), ‘intergenic’ (10.1%), and ‘UTR5’ (8.8%) being the most common. For VEP, 33,756 variants were discordant, with ‘downstream’ (41.0%), ‘upstream’ (21.8%), ‘nonsynonymous’ (13.5%), and ‘non-coding intron’ (7.6%) being the most common.

The row-normalized (*i.e*., normalized to total VEP annotations in each category), logtransformed consequence heatmap for VEP versus ANNOVAR on the Ensembl transcripts (Figure 2A) shows agreement for ‘inframe insertions’ and ‘inframe deletions’, ‘5’’ and ‘3’ untranslated region’, ‘intronic’, ‘upstream’, ‘downstream’, and ‘intergenic’ assignments. However, where PyVar reports ‘None’ (a null or missing value) from VEP, ANNOVAR often assigned ‘non-coding exonic.’ Annotations such as ‘stoploss’, ‘stopgain’, ‘frameshift’, ‘nonsynonymous’, and ‘synonymous’ showed some disagreement between the methods. For the RefSeq transcript set heatmap (Figure 3A), only ‘inframe insertion’ and ‘inframe deletion’ show strong concordance. Looking at the same data when column-normalized (normalized to total ANNOVAR annotations for each category; Figure 2B) shows concordance for ‘stoploss’, ‘stopgain’, ‘inframe insertion‘, ‘inframe deletion’ and ‘frameshift.’ ‘Stoploss’ calls by ANNOVAR are sometimes called as ‘frameshift’ by VEP. Similarly, both methods appear to disagree on ‘synonymous’ versus ‘nonsynonymous‘ calls.

**Figure 2.**
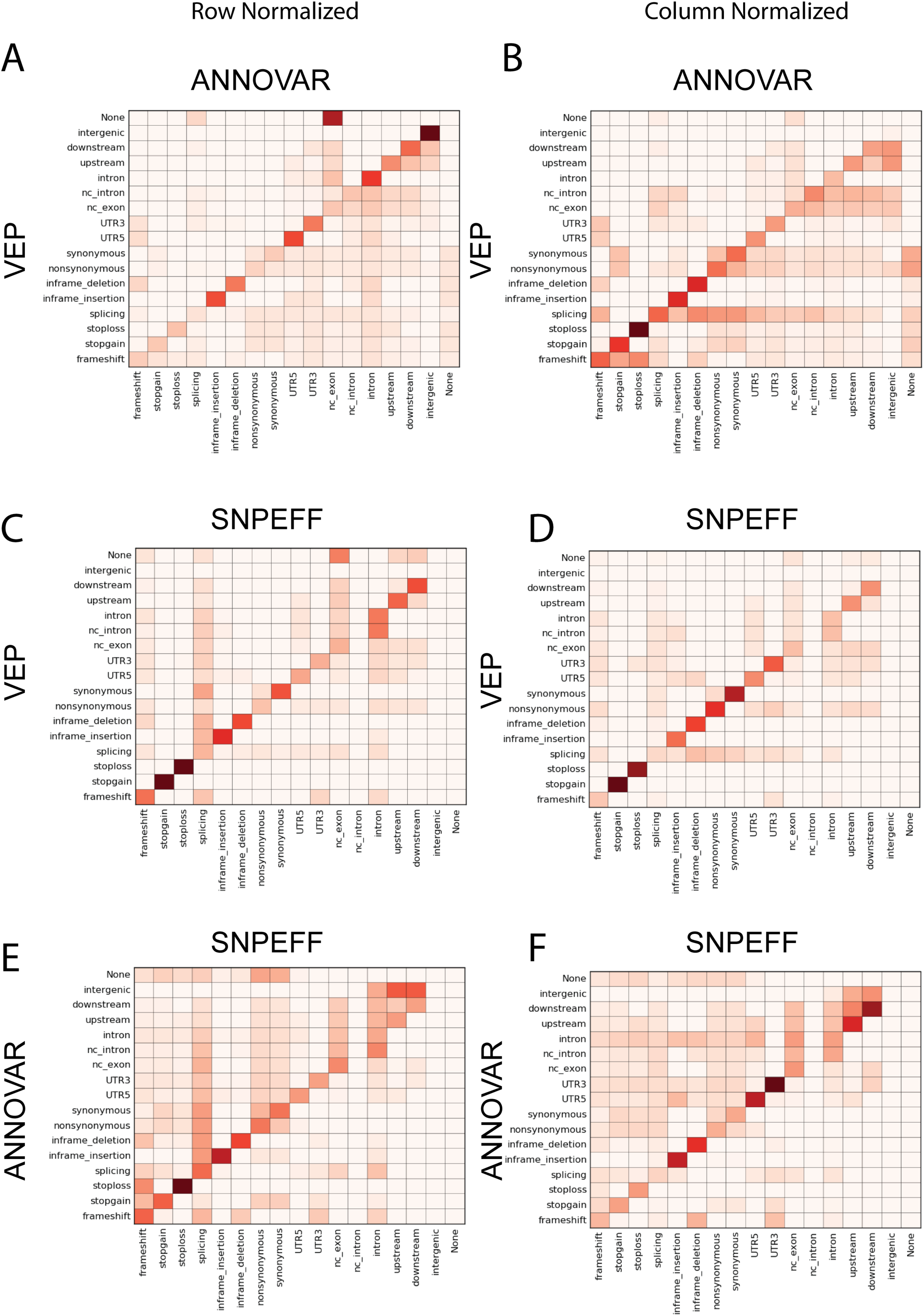
Normalized consequence heatmaps for Ensembl transcript set. Counts of the most severe normalized consequence identified by each annotator (color correspond to the log of the raw counts). Heatmaps are row-normalized (left) and column-normalized (right). **2A** and **2B** represent the comparison of consequences between VEP and ANNOVAR. **2C** and **2D** between VEP and SnpEff. **2E** and **2F** between ANNOVAR and SnpEff.

#### SnpEff versus VEP

When comparing the results of annotation by SnpEff and VEP, we found no position mismatches. With Ensembl transcripts, both methods used at least one common transcript for annotation > 99.9% of the time. However, for RefSeq transcripts, the concordance was lower with only 85.8% of variants sharing a common transcript for annotation. As before, ‘intronic’, ‘downstream’ and ‘upstream’ were among the most common annotations sharing a RefSeq transcript. Among the ~21,000 variants with a unique RefSeq transcript from VEP, the majority were annotated as ‘downstream’ (50.6%), ‘upstream’ (23.0%), ‘non-coding intronic’ (6.1%), and ‘non-synonymous’ (5.8%). Among the ~21,000 variants annotated with unique RefSeq transcripts by SnpEff, the majority were ‘UTR3’ (38.4%), followed by ‘intronic’ (27.9%), ‘intergenic’ (8.1%), and ‘UTR5’ (6.8%).

The row-normalized (VEP-normalized) consequence heatmap for VEP versus SnpEff using Ensembl (Figure 2C) shows concordance when VEP calls ‘frameshift’, ‘stopgain’, ‘stoploss’, ‘inframe insertion’, ‘inframe deletion’, ‘synonymous’, ‘intron’, ‘upstream’ and ‘downstream’. In some cases when SnpEff reported ‘splicing’, ‘downstream/upstream’, and ‘frameshift’, we noted that VEP had missing values (‘None’). When looking at the RefSeq row-normalized (VEP consequence-normalized) heatmap (Figure 3C), the concordance is not as strong. In particular, there is poorer agreement on ‘downstream’, ‘upstream’, ‘intron’, ‘non-coding intron/exon’, and ‘UTR5/3’. Even VEP calls of ‘synonymous’ and ‘non-synonymous’ show poor agreement with SnpEff calling ‘UTR3/UTR5’, ‘intron’, ‘upstream’ and other categories instead.

**Figure 3.**
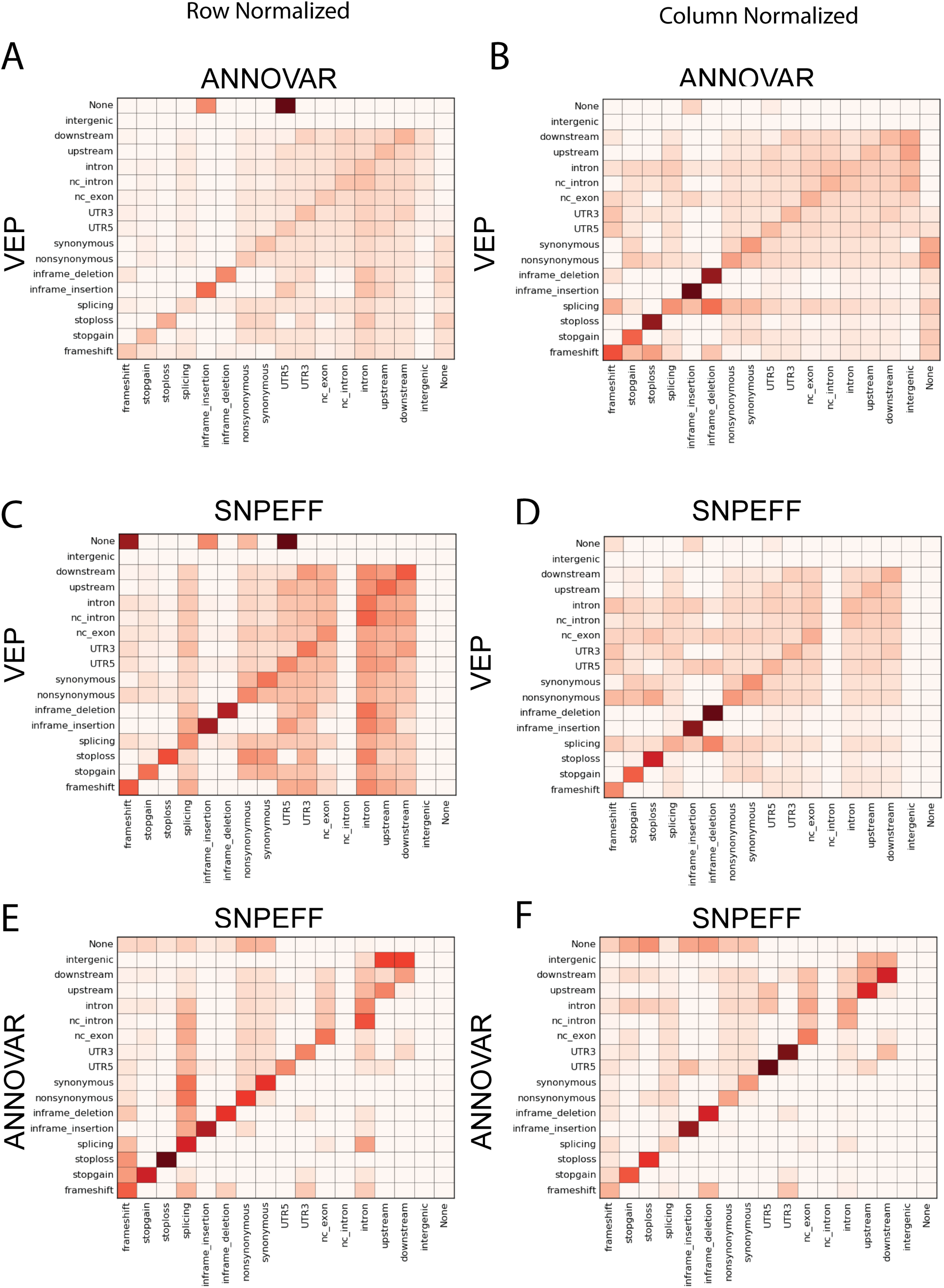
Normalized consequence heatmaps for the RefSeq transcript set. Counts of the most severe normalized consequence identified by each annotator (color correspond to the log of the raw counts). Heatmaps are row-normalized (left) and column-normalized (right). **3A** and **3B** represent the comparison of consequences between VEP and ANNOVAR. **3C** and **3D** between VEP and SnpEff. **3E** and **3F** between ANNOVAR and SnpEff.

The column-normalized (SnpEff-normalized) consequences heatmap for Ensembl transcripts (Fig 2D) shows good concordance for a majority of classifications, with the exception of ‘splicing’ and ‘frameshift.’ For RefSeq transcripts (Figure 3D), the heatmap shows that SnpEff’s call of ‘inframe deletion’ is sometimes called as ‘splicing’ by VEP. When SnpEff calls ‘splicing’ there is very little agreement with VEP. This applies to both RefSeq and Ensembl transcript sets. There is also poor agreement for ‘nonsynonymous’, ‘synonymous’, ‘UTR5’, ‘UTR3’, ‘intron’, ‘upstream’, and ‘downstream’ using RefSeq transcripts.

#### ANNOVAR versus SnpEff

ANNOVAR and SnpEff agreed on variant position in 99.7% of cases for both RefSeq and Ensembl transcript sets. For Ensembl transcripts, a common transcript was annotated for 61.5% of variants (Figure 1). For RefSeq, this was 50.2%. Of the ~52,000,000 variants sharing an Ensembl transcript between methods, ~44,700,000 also shared a common annotation, with ‘intron’ being the most abundant (76.7%). Of the 7,600,000 variants that had discordant annotations from SnpEff, nearly all (99.9%) were called as ‘intron.’ Similarly, the 7,600,000 variants with discordant calls from ANNOVAR were all called as ‘non-coding intron’. Of the 42,000,000 variants sharing at least one RefSeq transcript for annotation, 38,700,000 also shared a common annotation. 3,700,000 variants for SnpEff and ANNOVAR were annotated with unique consequences. As above, the shared annotations consisted mainly of ‘intron’ (82.6%). Where discordant annotations were made, they were ‘intron’ (99.7%) for SnpEff, and ‘non-coding intron’ (100%) for ANNOVAR.

The row-normalized heatmap for Ensembl transcripts (Figure 2E) shows good agreement for ‘frameshift’, ‘stoploss’, ‘inframe insertion’, ‘inframe deletion’, ‘UTR5’, and ‘UTR3’. The ‘splicing’ annotation has poorer agreement, as well as ‘synonymous’ and ‘nonsynonymous.’ The pattern is similar for row-normalized RefSeq consequences (Figure 3E), with good agreement along the diagonal for most annotations, with the exception of ‘splicing’. In the column-normalized (SnpEff-normalized) view (Figure 2F) of the heatmap data for Ensembl, we see that ANNOVAR gives ‘downstream’ and ‘intergenic’ annotations to SnpEff’s ‘upstream’ frequently. Similarly, what SnpEff calls ‘downstream’ is called as ‘intergenic’ by ANNOVAR. There is also poor agreement for ‘splicing’ and ‘frameshift’ calls.

### Concordance between methods for LoF variants for Ensembl transcripts

Approximately 653,000 variants were given loss-of-function annotations by either ANNOVAR or VEP (defined here as one or more of ‘nonsynonymous’, ‘stopgain’, ‘stoploss’, and ‘frameshift’; ‘splicing’ could also be LoF, however we chose not to include it in this analysis). This represents 0.8% of all variants analyzed. Of these 653,000 variants, just 934 (0.1%) were discordant between ANNOVAR and VEP, with annotators predicting different consequences. Figure 4 shows that the most common discordant annotation was ‘nonsynonymous’ (529), followed by multiple predicted consequences such as ‘nonsynonymous, downstream’ (85), and ‘nonsynonymous, upstream, downstream’ (68).

**Figure 4.**
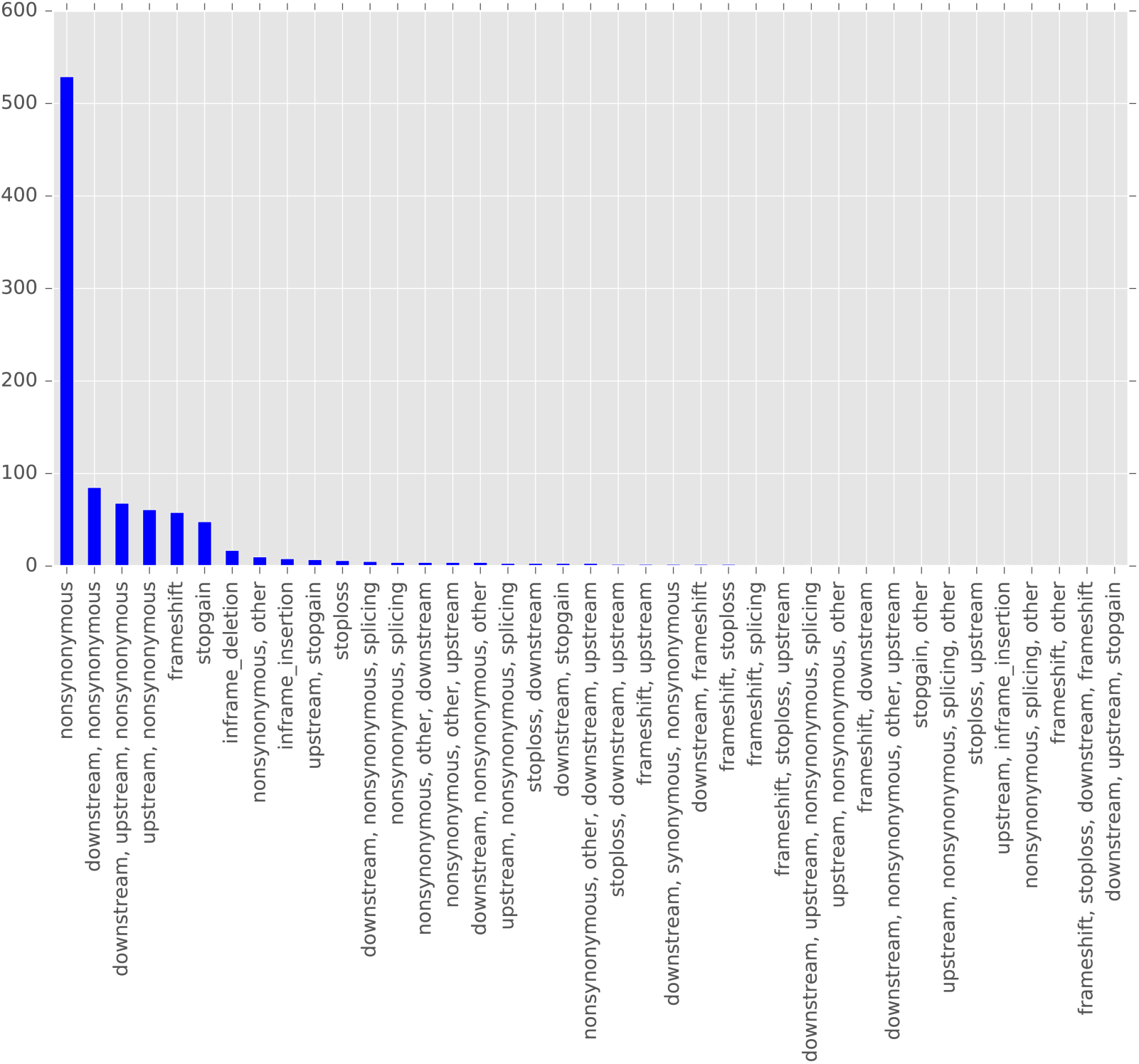
ANNOVAR versus VEP discordant annotations (on LoF variants).

When comparing ANNOVAR and SnpEff (Figure 5), ~654,000 variants were annotated as LoF by one or both methods. Of those, 1456 (0.2%) were discordant between the methods. ‘Nonsynonymous’ alone or in combination with other annotations accounted for the six most common discordant annotations. Between SnpEff and VEP, there were 63 discordant annotations among only ~456,500 variants called as LoF (0.01%). ‘Nonsynonymous, other, frameshift’ (23) and ‘nonsynonymous, frameshift’ (12) were the top two most common discordant annotations between methods (Figure 6).

**Figure 5.**
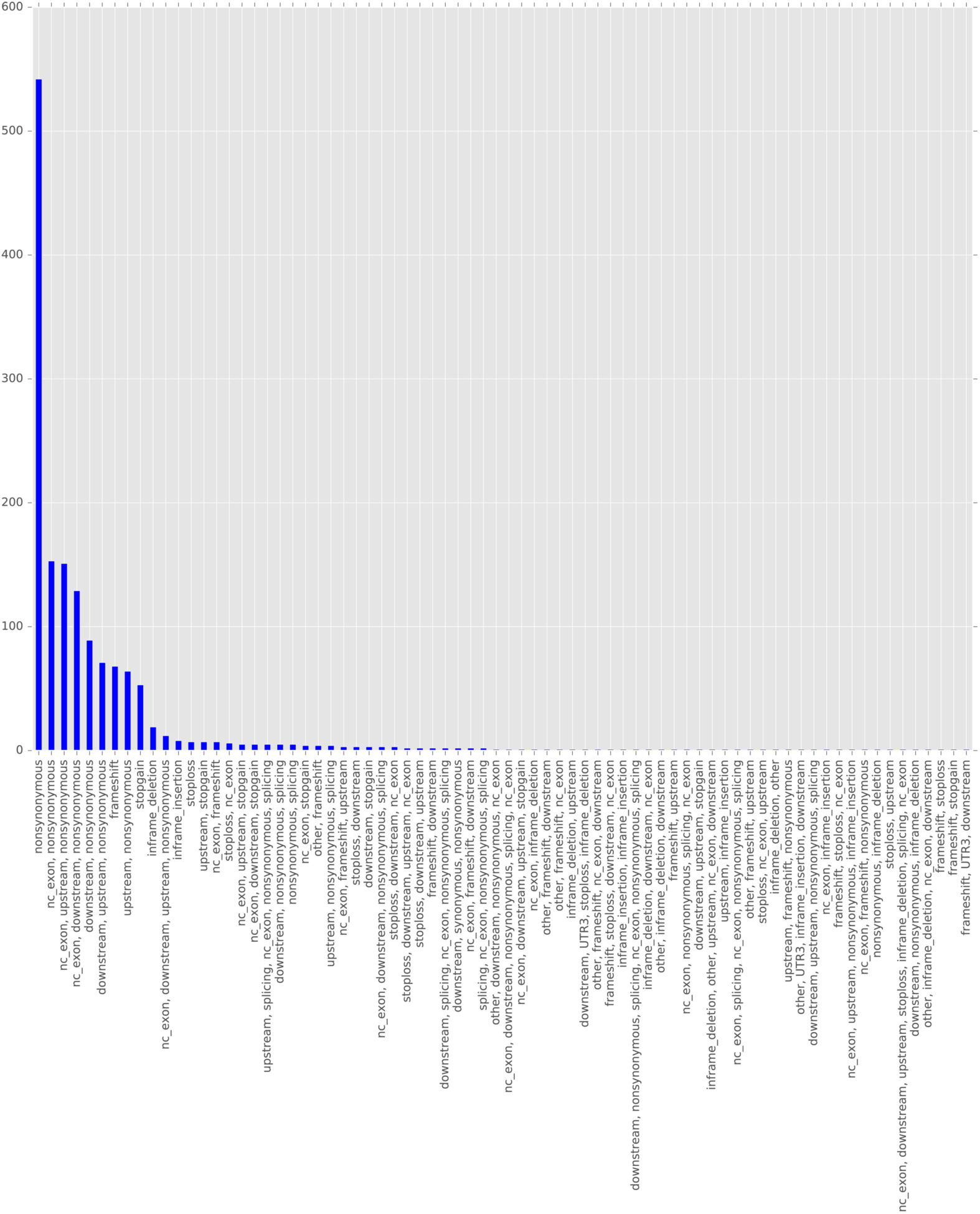
ANNOVAR versus SnpEff discordant annotations on LoF variants.

**Figure 6.**
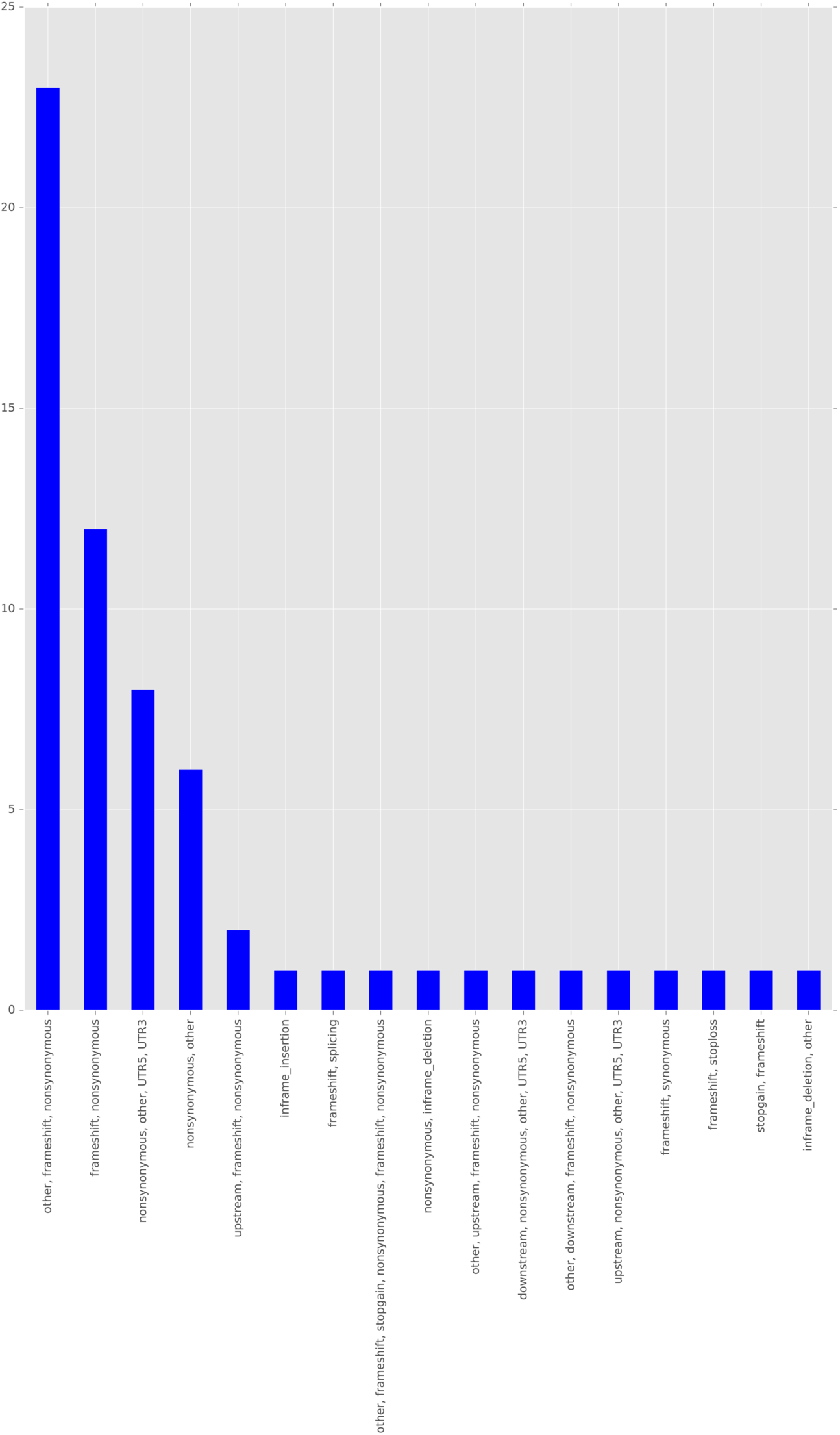
SnpEff versus VEP discordant annotations on LoF variants.

## Discussion

The small amount (< 0.5%) of disagreement between methods on variant position, for example, between ANNOVAR and VEP using the RefSeq transcripts can be attributed to different approaches between methods for assigning numbering schemes to insertions and deletions. The majority of these variants are found in intronic and intergenic regions and are generally not high-priority LoF variants. It is important to be aware, however, that the methods may not agree on variant start position in the case of certain indels.

One unexpected discrepancy that emerges when comparing ANNOVAR, VEP, and SnpEff is the lack of concordance for a very high percentage of variants when the annotator methods are choosing transcripts. In some comparisons, this was as high as 55%. Thus, a large number of variants, the vast majority of which are intergenic, appear to be annotated on different transcripts. Looking more closely at the output from each method shows that these discordant transcript choices arise from the classification of a variant in an intergenic region as belonging to an ‘unknown’ transcript (VEP) or belonging to the nearest transcript in the sequence (ANNOVAR). Although this is mainly affecting variants that are typically not LoF and therefore not high-priority, it is important to be aware that methods disagree substantially on this basis. In the case of what appears to be 100% agreement on transcript choice between SnpEff and VEP, this is owing to the use of the ‘unknown’ designation by both methods, rather than attempting, as ANNOVAR does, to assign a transcript based on proximity.

The heatmaps of normalized consequences for the three methods show that overall, concordance is better with Ensembl rather than RefSeq transcript sets. Disagreement on assigning a call of ‘splicing’ to a variant is a common thread between all methods, both with Ensembl and RefSeq. This is not surprising given that prediction of splicing defects is a challenging problem that remains open to further research and validation [7,8]. Calls of ‘inframe insertions’ and ‘deletions’, ‘nonsynonmous’ and ‘synonymous’, ‘stopgain’, ‘stoploss’, and ‘frameshift’ all showed the best concordance between methods, particularly when using Ensembl transcripts. Assignments of ‘intergenic,’ ‘downstream,’ ‘upstream,’ and ‘intron’ were a source of disagreement between methods regardless of Ensembl or RefSeq transcripts. These disagreements could possibly be alleviated by adjusting parameters within the methods to redefine the boundary between up- and downstream and intergenic regions (although we did not test this). In general, concordance between methods was poorer when using RefSeq transcripts for reasons that are not well understood. RefSeq contains fewer transcripts by a significant percentage than Ensembl, and generally has less complex gene models [9]. Since concordance is defined as at least one consequence in common, the greater number of transcript choices in Ensembl could be responsible for the greater concordance seen for that transcript set.

The heatmaps in Figures 1 and 2 are row- and column-normalized and log-transformed to visually emphasize the rare discordant annotations between methods. However, our analysis of just LoF variants shows that discordance for this clinically-important category of variants is, in our hands, very low. Others have found much higher rates of discordance, as high as 36% [2], (this figure includes splicing and indels which we are excluding). Excluding splicing and indel variants, the authors in that study find ~14% discordance. This is still substantially larger than the rate of 0.1% between ANNOVAR and VEP that we detect here and could be owing to differences in command-line parameters and filtering of variants. Additionally, in that work the authors studied variants from 276 genomes from individuals with immune disease, Mendelian disease, and cancer. Our dataset, in contrast, comes from the 1000 Genomes project and is not biased for individuals with genetic diseases. It is possible that differences in the kinds of variants found in the underlying variant dataset are responsible for these discrepancies.

## Conclusions

In conclusion, we have created a python framework, PyVar, for automatically comparing the results of variant annotation from different popular methods and for reconciling different output formats and consequence ontologies to allow easy exploratory analysis. We tested our method on ~80 million variants from the 1000 Genomes Project and showed that there was significant disagreement on transcript choice between methods owing to each method’s procedures for handling intergenic and intronic variants. However, for more interesting LoF variants (excluding splicing), we found that concordance was actually very good. It is our hope that the community can find value in our PyVar framework for quick automated comparisons between novel and existing methods.

## Methods

### Data acquisition

The data used in this paper was obtained from the publically available FTP repository at the 1000 Genomes Project: ftp://ftp.1000genomes.ebi.ac.uk/vol1/ftp/release/20130502/ \ ALL.wgs.phase3_shapeit2_mvncall_integrated_v5b.20130502.sites.vcf.gz. The variant calls are from the phase3 release of May 2, 2013. The variant set contains calls from 2504 individuals from 26 populations. There are 78,136,341 SNPs and 3,135,424 indels in the VCF file.

### Variant annotation

Variant annotations were obtained using ANNOVAR 2016.2.1, VEP ver. 83, and SnpEff ver. 4.2. The programs were invoked with the following commands:

#### SnpEff/Ensembl

java -Xmx16g -jar /Users/wrtz/SnpEff/SnpEff.jar GRCh37.75

ALL.wgs.phase3_shapeit2_mvncall_integrated_v5b.20130502.sites.vcf >

/Users/wrtz/annotation_paper/ALL.wgs.phase3_fixed.vcf.SnpEff_ens_out

#### SnpEff/RefSeq

java -Xmx16g -jar /Users/wrtz/SnpEff/SnpEff.jar hg19

ALL.wgs.phase3_shapeit2_mvncall_integrated_v5b.20130502.sites.vcf >

/Users/wrtz/annotation_paper/ALL.wgs.phase3_fixed.vcf.SnpEff_refseq_out

#### VEP/Ensembl

perl /Users/wrtz/vep3/ensembl-tools-release-84/scripts/variant_effect_predictor/variant_effect_predictor.pl --port 3337 --cache --assembly GRCh37 -i ALL.wgs.phase3_shapeit2_mvncall_integrated_v5b.20130502.sites.vcf -o ALL.wgs.phase3_shapeit2_mvncall_integrated_v5b.20130502.sites.ens.vep.vcf --fork 8 --vcf

#### VEP/RefSeq

perl /Users/wrtz/vep3/ensembl-tools-release-84/scripts/variant_effect_predictor/variant_effect_predictor.pl --port 3337 --cache --assembly GRCh37 -i ALL.wgs.phase3_shapeit2_mvncall_integrated_v5b.20130502.sites.vcf -o ALL.wgs.phase3_shapeit2_mvncall_integrated_v5b.20130502.sites.refseq.vep.vcf --fork 8 --refseq --vcf

#### ANNOVAR/Ensembl

perl convert2annovar.pl -includeinfo -allsample -withfreq -format vcf4 ALL.wgs.phase3_fixed.vcf > ALL.wgs.phase3.avinput;

perl annotate_variation.pl -out ALL.wgs.phase3_shapeit2_mvncall_integrated_v5b.20130502.sites.ens.annovar - geneanno -buildver hg19 -separate -neargene 5000 -transcript_function -hgvs -splicing_threshold 5 -thread 8

../ALL.wgs.phase3.avinput humandb/ -dbtype ensGene

#### ANNOVAR/RefSeq

perl convert2annovar.pl -includeinfo -allsample -withfreq -format vcf4 ALL.wgs.phase3_fixed.vcf > ALL.wgs.phase3.avinput;

perl annotate_variation.pl -out ALL.wgs.phase3_shapeit2_mvncall_integrated_v5b.20130502.sites.refseq.annovar - geneanno -buildver hg19 -separate -neargene 5000 -transcript_function -hgvs -splicing_threshold 5 -thread 8 ../ALL.wgs.phase3.avinput humandb/

#### PyVar framework

The PyVar framework was written in python 2.7 (http://www.python.org) and requires the following additional libraries: matplotlib, matplotlib_venn, and numpy. It can be downloaded from the PyVar GitHub repository (https://github.com/jwertz01/annotator-comparison).

#### Variant annotation comparison

By default, PyVar converts the consequences from each annotator method into a standardized nomenclature for useful comparison. Table 1 summarizes the conversions between the output nomenclature of each method and the standardized terms used by PyVar. PyVar retained only variants sharing a common position for the next step (analysis of transcript commonality). Similarly, PyVar retained only those variants sharing at least one transcript between methods for heatmap analysis of normalized consequences.

Examples of the command-line code used to invoke PyVar are shown below:

ANNOVAR versus SnpEff (w/ RefSeq transcripts)

python ../compare_annotators.py --anv_var_func_filenames

../annovar/ALL.wgs.phase3_shapeit2_mvncall_integrated_v5b.20130502.sites.refseq.annovar.variant_function -- anv_exonic_var_func_filenames

../annovar/ALL.wgs.phase3_shapeit2_mvncall_integrated_v5b.20130502.sites.refseq.annovar.exonic_variant_functio n --SnpEff_vcf_filenames ../ALL.wgs.phase3_fixed.vcf.SnpEff_refseq_out --filegroup_1

ALL.wgs.phase3_shapeit2_mvncall_integrated_v5b.20130502.sites.refseq.annovar.variant_function

ALL.wgs.phase3_shapeit2_mvncall_integrated_v5b.20130502.sites.refseq.annovar.exonic_variant_function

#### SnpEff versus VEP (w/ Ensembl transcripts)

python ../compare_annotators.py --vep_vcf_filenames

../ALL.wgs.phase3_shapeit2_mvncall_integrated_v5b.20130502.sites.ens.vep.vcf --SnpEff_vcf_filenames

../ALL.wgs.phase3_fixed.vcf.SnpEff_ens_out

#### ANNOVAR versus VEP (w/ Ensembl transcripts)

python ../compare_annotators.py --vep_vcf_filenames

../ALL.wgs.phase3_shapeit2_mvncall_integrated_v5b.20130502.sites.ens.vep.vcf --anv_var_func_filenames

../annovar/ALL.wgs.phase3_shapeit2_mvncall_integrated_v5b.20130502.sites.ens.annovar.variant_function -- anv_exonic_var_func_filenames

../annovar/ALL.wgs.phase3_shapeit2_mvncall_integrated_v5b.20130502.sites.ens.annovar.exonic_variant_function --filegroup_1 ALL.wgs.phase3_shapeit2_mvncall_integrated_v5b.20130502.sites.ens.annovar.variant_function

ALL.wgs.phase3_shapeit2_mvncall_integrated_v5b.20130502.sites.ens.annovar.exonic_variant_function

## Author contributions

JW, TB, and MSC conceived the experiments. JW wrote the PyVar code framework. JW and MSC conducted the analysis. MSC wrote the manuscript.

